# A transcriptional signature of LKB1 functional loss defines a large, therapeutically addressable patient population across human cancers

**DOI:** 10.64898/2026.07.23.740219

**Authors:** Sourav Bandyopadhyay, John D. Gordan

## Abstract

Loss of the tumor suppressor LKB1 (STK11) drives metabolic reprogramming and immune evasion, but its clinical footprint is defined almost entirely by somatic mutation and deletion. Because emerging therapies aim to reactivate this pathway in structurally wild-type tumors, defining the full LKB1-deficient population (including tumors silenced by non-genomic mechanisms) is a prerequisite for patient selection. We derived a 30-gene transcriptional signature of LKB1 functional loss, training on genomically-defined STK11 loss in lung adenocarcinoma and validating in an independent squamous cohort (AUROC 0.926). It transferred to four independent non-TCGA LUAD cohorts (AUROC 0.92–0.98), outperforming the published Kaufman 16-gene classifier. Critically, restoring wild-type LKB1 in LKB1-mutant NSCLC lines reversed the signature while a kinase-dead mutant did not, establishing that it reads out LKB1 kinase function, not merely mutation status. Applied pan-cancer, the signature identified functional LKB1 loss in wild-type tumors at 7.2%, expanding total LKB1-loss prevalence 3.7-fold over genomic loss alone (2.7% → 9.9%), largest in esophageal, colorectal, endometrial and cutaneous cancers. This population is decoupled from LKB1 mutation and deletion frequency, yet enriched in BRAF-mutant colorectal (25% signature-positive) and HER2-amplified breast (21% signature-positive) cancers: readily-testable subgroups for screening. In lung adenocarcinoma, signature positivity rose across a co-mutation gradient from 6% background to 15% in KRAS-mutant, 35% in NRF2-pathway-mutant (KEAP1 or NFE2L2), to 64% in KRAS/NRF2 co-mutant tumors. Similar to KRAS/LKB1 co-mutant tumors, LKB1 wild-type but signature-high tumors were immune-cold, correlating negatively with antigen-presentation, interferon-γ and T-cell-inflamed programs in multiple independent lung cancer cohorts. These results define a broad set of tumors with LKB1 functional loss that may open up new avenues for precision therapeutics.

## Introduction

LKB1, encoded by STK11, is a serine/threonine kinase and one of the most broadly acting tumor suppressors in human cancer. It was identified through the germline loss-of-function mutations that cause Peutz-Jeghers syndrome, an autosomal-dominant polyposis with markedly elevated lifetime risk of gastrointestinal, pancreatic, breast, lung and gynecological cancers [1,2,3]. LKB1 exerts its tumor-suppressor activity as a heterotrimeric complex with the pseudokinase STRAD and the scaffold MO25, which localizes and activates the kinase in the cytoplasm [4]. In its active form LKB1 is the principal upstream kinase of AMP-activated protein kinase (AMPK) [5,6,7], coupling LKB1 to cellular energy sensing and, through AMPK-dependent restraint of mTORC1, to the control of anabolic growth [8]. LKB1 additionally activates a family of thirteen AMPK-related kinases that govern chromatin accessibility, cell polarity and cytoskeletal organization [9], so that LKB1 loss simultaneously deregulates metabolism, growth signaling and epithelial architecture [10,11,12,13].

The clinical consequences of LKB1 loss are best characterized in lung adenocarcinoma (LUAD), where STK11 is among the most frequently inactivated tumor suppressors and its loss is enriched in adenocarcinoma histology and in smokers [14]. In genetically engineered models, Lkb1 deletion accelerates Kras-driven lung tumorigenesis and promotes metastasis and histological plasticity [15]. Co-mutation of STK11 with KRAS defines a distinct molecular subset (the “KL” subgroup) with an immune-cold microenvironment [16], driven mechanistically by suppression of STING-dependent innate immune sensing and by neutrophil-mediated T-cell suppression [17,18,19]. This subset shows primary resistance to PD-1 blockade [20,21], and LKB1 loss more broadly exposes metabolic vulnerabilities that have been proposed as therapeutic entry points [22]. Recent work has defined a novel small-molecule approach for activating LKB1 in LKB1-wild-type tumors, raising the possibility of reversing the tumorigenic consequences of losing this tumor suppressor [48].

A central limitation of the existing prevalence literature is that it is defined largely by mutation. LKB1 can be inactivated without a coding mutation through several orthogonal routes. Promoter CpG-island hypermethylation silences LKB1 in colorectal and cervical carcinoma, where somatic mutation is comparatively rare [23,24], and non-mutational loss of function has been tied to global DNA-methylation reprogramming in LUAD [25,26]. Transcriptional down-modulation via an NKX2-1/p53 axis produces functional LKB1 loss in colorectal cancer in the absence of mutation, LOH or methylation [27]. Post-transcriptional silencing by microRNAs (miR-17, miR-100) identifies STK11-wild-type tumors that are nonetheless LKB1-deficient [28,29], and post-translational mechanisms can eliminate LKB1 protein despite an intact, expressed gene [30,31]. Consistent with this, immunohistochemical loss of LKB1 protein marks aggressive biology in KRAS-mutant LUAD independently of mutation status [32]. The same theme recurs across cervical, pancreatic and biliary neoplasia [33,34,35,36]. Because these mechanisms are invisible to mutation calling, the true LKB1-deficient population is larger than sequencing-based estimates imply, and its size per cancer type is poorly quantified [37].

Transcriptional signatures offer a mechanism-agnostic readout of LKB1 loss because they capture the downstream consequences of pathway inactivation regardless of cause. The Kaufman 16-gene classifier, derived and applied in NSCLC, predicts LKB1 inactivation by both mutational and non-mutational mechanisms and is attenuated when wild-type LKB1 is restored [38]. Other definitions are tumor-type-specific [16] or mouse-derived [39], and the reported divergence between human and murine LKB1-loss signatures [38] cautions against transferring model-derived definitions to human tumors.

Whether a single expression-based definition transfers across human tumor types, tracks LKB1 kinase function causally, and can quantify the functionally-deficient population pan-cancer has not been established. Here we address these gaps: we derive a transcriptional signature from genomically-defined loss in TCGA, validate it across independent cohorts, cell lines and an LKB1 add-back experiment, and use it to quantify functional LKB1 loss per cancer type and to map its genomic and immunological context.

## Methods

### Data source and cohorts

We analyzed the 32 TCGA PanCancer Atlas studies available through cBioPortal (10,443 sequenced tumors), retrieving STK11 somatic mutation and copy-number data, RNA-seq expression (RSEM), RPPA LKB1 protein, and hm450 promoter-CpG beta values. Independent validation used three non-TCGA LUAD cohorts with paired expression and STK11 mutation data (OncoSG [40], CPTAC [41], CAS [42]) and the SU2C-MARK immunotherapy cohort (152 NSCLC tumors with harmonized RNA-seq and 309 with whole-exome sequencing) [46]. Cell-line data (expression, STK11 mutation/copy-number, CRISPR dependency, drug sensitivity) were from the Cancer Cell Line Encyclopedia and DepMap [47]. LKB1 add-back data were the isogenic A549 and H2122 series from Kaufman et al. (GSE51266): each LKB1-mutant line transduced with empty vector, wild-type LKB1, or kinase-dead LKB1-K78I, in triplicate [38]. Instructions and code to reproduce the findings in this manuscript can be accessed at: https://github.com/souravUCSF/lkb1-functional-loss

### Genomic LKB1 loss

Deleterious STK11 events (nonsense, frameshift, splice-site, translation-start, other truncating) or homozygous deletion (GISTIC = −2) defined genomic loss; tumors with neither were structurally wild-type (WT).

### Signature derivation

Expression was log2(RSEM+1)-transformed. The signature was derived in LUAD (largest genomic-loss set): positives = genomic loss (n = 55); negatives = confidently-intact controls (WT with STK11 expression ≥ cohort 40th percentile, n = 177). An elastic-net-penalized logistic regression with class weighting was tuned in nested cross-validation (outer 5-fold for unbiased AUROC/AP; inner 4-fold for feature selection and penalty hyperparameters), then locked by refitting on the full discovery set. Non-zero-coefficient genes constitute the 30-gene signature.

### Validation and benchmarking

The locked signature was applied to LUSC using discovery reference statistics. Cross-tumor generalization was assessed by within-cohort standardization in every cohort with ≥5 genomic-loss cases. External cohorts (OncoSG, CPTAC, CAS, SU2C-MARK) were standardized within-cohort and scored with the locked weights; discrimination of STK11 loss was measured by AUROC. The signature was benchmarked against the Kaufman 16-gene classifier (mean z-score of constituent genes) across all six cohorts. In cell lines, discrimination of genomic STK11 loss was measured overall and within the lung subset. In the add-back experiment, the signature was scored within each cell line (within-line z of the 28/30 genes present on the array) and compared across conditions by t-test.

### Functional-loss calling

In structurally-WT tumors, per-cohort signature-positive thresholds were set at 95% specificity in confidently-intact controls (5% false-positive floor); functional-loss estimate = excess positivity above this floor, with beta-distribution intervals. Cohorts were signature-validated when they carried ≥5 genomic-loss cases.

### Genomic correlates

Across a 30-gene panel of clinically-assessed cancer genes in WT tumors, associations with functional-loss status were tested by cohort-stratified Cochran-Mantel-Haenszel statistics and multivariable logistic regression with tumor type as a fixed effect (Benjamini-Hochberg-corrected); clinical co-occurrences were tested within tumor types by Fisher’s exact test.

### NSCLC co-mutation and NRF2 axis

In TCGA LUAD, WT tumors were stratified by KRAS mutation and NRF2-pathway mutation (KEAP1 or NFE2L2). Signature positivity was tabulated across the four strata and the trend tested by Spearman correlation; KEAP1/NRF2-pathway co-occurrence was decomposed by LKB1 status (genomic loss vs functional-loss-WT vs signature-negative-WT).

### Immune-pathway analysis

Per-sample immune scores (mean within-cohort z of published gene sets: IFN-γ, T-cell-inflamed GEP, PD-1/PD-L1 axis, cytolytic activity, antigen presentation) were correlated with the signature score in TCGA LUAD (n = 510) and SU2C-MARK (n = 152) by Spearman correlation, both across all tumors and within genomically-WT tumors alone.

## Results

### A 30-gene signature reads out LKB1 functional loss and is common across cancers

Across the 32 TCGA tumor types, genomic STK11/LKB1 loss was strongly concentrated in LUAD (10.1%), followed by cervical (5.8%) and low single-digit frequencies elsewhere, confirming that mutation-based ascertainment identifies few LKB1-deficient tumors outside a handful of indications. In LUAD, genomic-loss tumors had markedly lower STK11 expression than WT tumors (median 8.86 vs 10.03 log2 RSEM; Mann-Whitney p = 1.7×10^-18^), justifying an expression-based classifier. Training penalized logistic regression on genomically-defined loss versus confidently-intact controls, with nested cross-validation, yielded a 30-gene signature (23 up, 7 down in loss) that discriminated genomic loss with an out-of-fold AUROC of 0.986 (average precision 0.975 against a 0.24 baseline) and 0.926 in an independent LUSC cohort (Figure 1a). The signature independently recovered SIK1 (a direct LKB1 substrate) and PDE4D from the Kaufman classifier without access to that gene list, both up-regulated in LKB1 loss (elastic-net weights +0.11 and +0.17); all 30 signature genes, with weights, directions and effect sizes, are shown in Figure S1. It retained discrimination with STK11 removed (mean within-tissue AUROC 0.68, up to 0.89 in LUAD and gastric cancer), showing it captures a downstream functional program rather than STK11 transcription alone. Promoter hypermethylation did not account for the low STK11 expression in genomic-loss tumors: the STK11 promoter CpG probes were constitutively methylated with no methylation–expression anticorrelation (ρ = −0.02, n = 452), confirming that a transcriptional rather than epigenetic readout is required (Figure S2). Displayed as a heatmap across the LUAD discovery cohort, the coordinated up- and down-regulation of the signature genes is evident in tumors with genomic LKB1 loss and in the structurally-wild-type tumors the signature calls functionally-deficient, while signature-negative wild-type tumors show the reciprocal pattern (Figure 1b).

**Figure 1.**
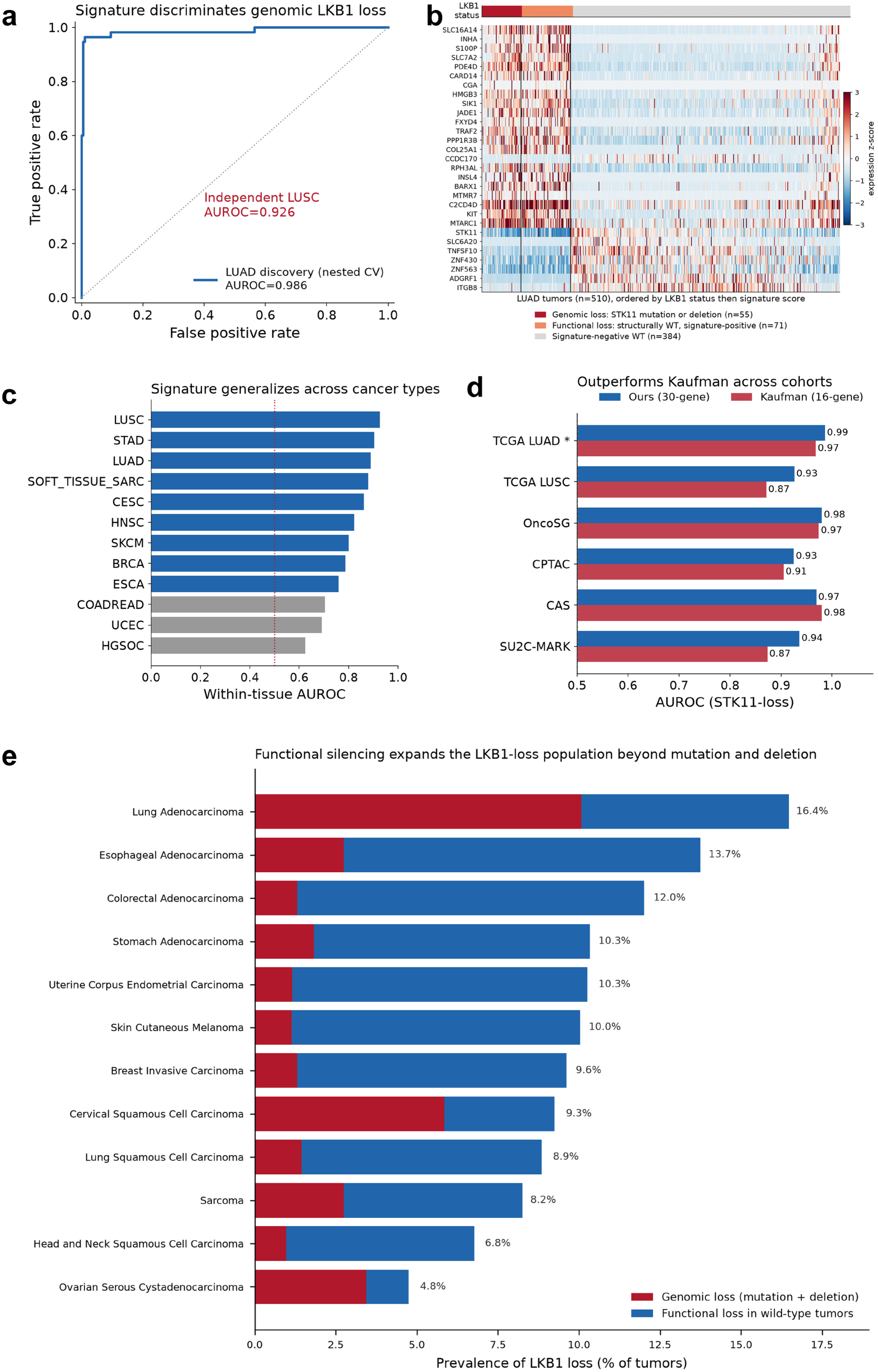
A 30-gene signature reads out LKB1 functional loss and is common across cancers. (a) ROC for discrimination of genomic LKB1 loss in the LUAD discovery cohort (out-of-fold AUROC 0.986) and independent LUSC (0.926). (b) Expression heatmap of the 29 signature genes measurable on the profile (z-scored) across 510 LUAD tumors, ordered by LKB1 status then signature score: genomic loss (deleterious STK11 mutation or homozygous deletion, n = 55), functional loss (structurally wild-type but signature-positive, n = 71) and signature-negative wild-type (n= 384). (c) Within-tissue AUROC across 12 cancers (mean 0.80); bars below 0.75 in grey. (d) Head-to-head against the Kaufman 16-gene classifier across six cohorts: TCGA LUAD (discovery; in-sample for our signature, independent for Kaufman) and LUSC plus four independent cohorts, with our signature winning 5/6 (pooled 0.95 vs 0.93). (e) Pan-cancer prevalence of LKB1 loss: genomic loss (red) plus functional loss in wild-type tumors (blue), showing that functional silencing expands the deficient population beyond mutation and deletion.

Standardized within each cohort, the signature generalized across tumor types (mean within-tissue AUROC 0.80 across 12 cancers), strongest in lung and gastric cohorts (Figure 1c). Benchmarked against the Kaufman 16-gene classifier across all six cohorts with genomic-loss cases: TCGA LUAD and LUSC plus the four independent cohorts our signature was superior in five of six (pooled AUROC 0.95 vs 0.93), remaining concordant with the Kaufman score (Spearman ρ = 0.71) while outperforming it (Figure 1d). Applying the validated signature to structurally-WT tumors revealed that functional loss substantially expands the deficient population beyond the STK11 mutation rate in most cancer types (Figure 1e).

### The signature transfers across cohorts and tracks LKB1 kinase function

To establish that the signature is not TCGA-specific, we applied it to four independent LUAD cohorts. Discrimination of STK11 loss was preserved in every one: OncoSG 0.980 (n = 169), CPTAC 0.925 (n = 110), CAS 0.969 (n = 51), and the SU2C-MARK immunotherapy cohort 0.936 (n = 68 matched RNA+WES, 13 genomic-loss cases) (Figure 2a–b). These closely match the TCGA discovery (0.986) and LUSC (0.926) estimates, establishing transfer across patient populations, sequencing platforms and geography. The signature also discriminated genomic STK11 loss in cancer cell lines (CCLE: AUROC 0.813 overall, 0.881 in the lung subset), extending validity to an orthogonal model system (Figure 2c–d).

**Figure 2.**
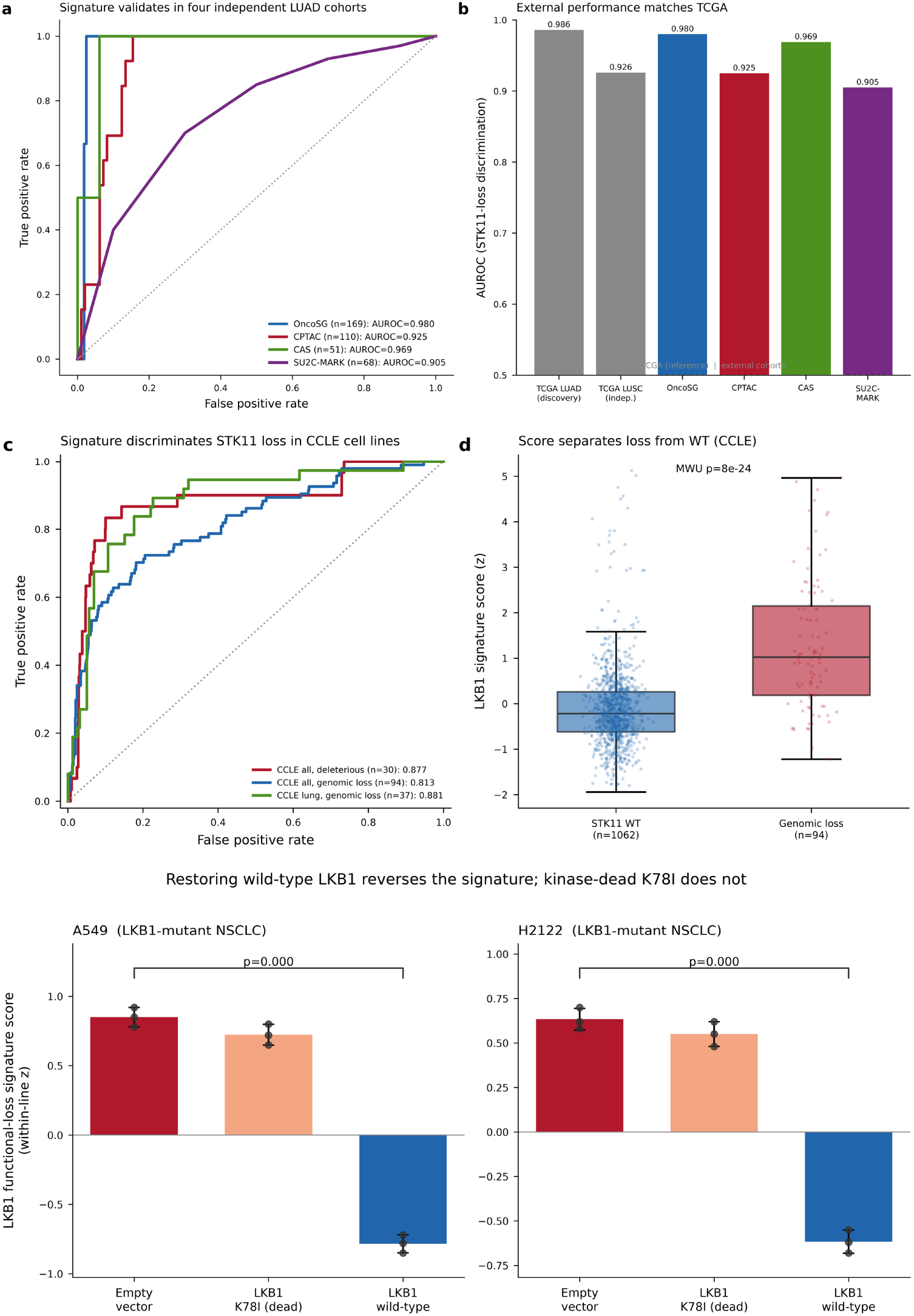
The signature transfers across cohorts and tracks LKB1 kinase function. (a) ROC and (b) AUROC bars for discrimination of STK11 loss in four independent non-TCGA LUAD cohorts: OncoSG (n = 169), CPTAC (n = 110), CAS (n = 51) and SU2C-MARK (n = 68), against the TCGA reference. (c) ROC and (d) score separation in CCLE cell lines (AUROC 0.813 overall, 0.881 in lung). (e) LKB1 add-back: signature score in isogenic A549 and H2122 LKB1-mutant NSCLC lines transduced with empty vector, kinase-dead LKB1-K78I, or wild-type LKB1. Wild-type LKB1 reverses the signature (p = 0.045, 0.016); kinase-dead K78I does not.

We then tested the therapeutic premise directly. In an independent isogenic add-back dataset (GSE51266; A549 and H2122 LKB1-mutant NSCLC lines transduced with empty vector, wild-type, or kinase-dead LKB1), re-expressing wild-type LKB1 significantly reversed the signature (A549 +0.85 → −1.38, p = 0.045; H2122 +3.21 → −2.38, p = 0.016), whereas the kinase-dead K78I mutant did not (A549 Δ = −0.32, n.s.; wild-type vs K78I H2122 p = 0.027) (Figure 2e). Signature reversal therefore requires LKB1 kinase activity, demonstrating that the signature reads out LKB1 pathway function rather than a correlate of mutation status.

### Functional LKB1 loss defines a large, structurally-intact target population

Across the 12 signature-validated cohorts (5,809 tumors), functional LKB1 loss in structurally-WT tumors affected 7.2% of all tumors (11.9% of WT), approximately 3.7-fold more patients than genomic loss alone (2.7%), raising total LKB1-loss prevalence to 9.9% (Figure 3a). The additional, functionally-deficient population was concentrated where STK11 mutation is uncommon: esophageal (11.0%), colorectal (10.7%), endometrial (9.1%), melanoma (8.9%), gastric (8.5%) and breast (8.3%) carcinomas. Plotting functional against genomic loss placed these cancers far above the diagonal (little mutational loss, substantial functional loss), with LUAD the converse (Figure 3b). The genomic component of this ranking was recapitulated in cell lines: CCLE STK11-loss prevalence by lineage tracked the TCGA tumor ranking (Pearson r = 0.73), led by non-small-cell lung cancer (Figure S3). Functional loss was likewise decoupled from STK11 mutation frequency across cancers (Figure 3c), indicating that multiple tumors types downregulate the LKB1 pathway though indirect means, most often more frequently than via mutation.

**Figure 3.**
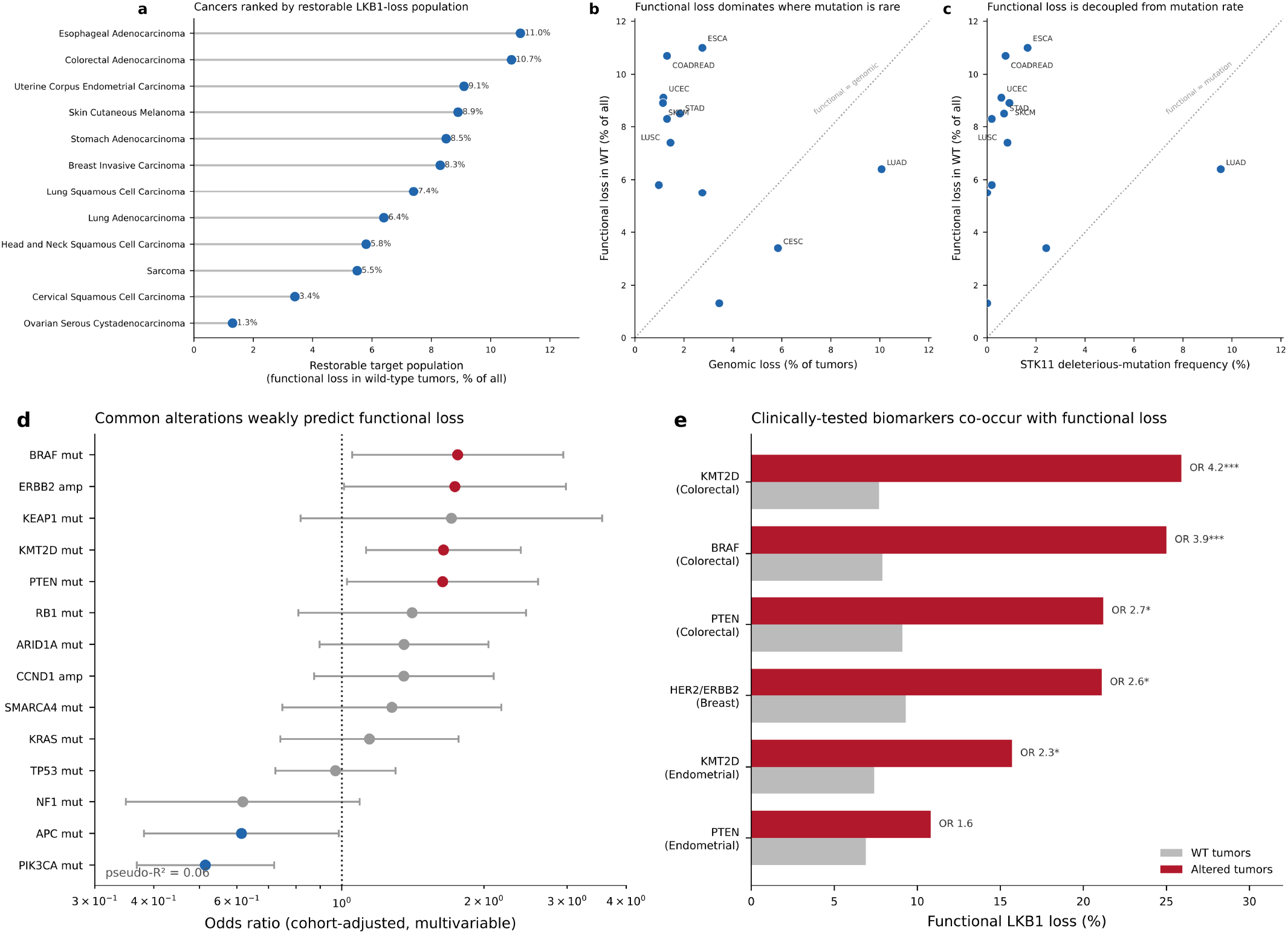
Functional LKB1 loss defines a large, structurally-intact target population. (a) Per-lineage prevalence of functional LKB1 loss in structurally wild-type tumors, ranked. (b) Functional versus genomic loss per cancer type; cancers above the diagonal carry little mutational but substantial functional loss. (c) Functional loss versus STK11 mutation frequency across cancers, showing decoupling. (d) Cohort-adjusted multivariable odds ratios for common alterations predicting functional loss in wild-type tumors (pseudo-R^2^ = 0.06), showing that the common driver landscape explains little of the phenotype. (e) Functional-loss rate in altered versus wild-type tumors for clinically-tested biomarkers, including BRAF-mutant colorectal (OR 3.9) and HER2-amplified breast cancer (OR 2.6). *p<0.05, **p<0.01**, *p<0.001.

A cohort-adjusted multivariable model across a 30-gene panel of clinically-assessed cancer genes explained little of the functional-loss phenotype in WT tumors (pseudo-R^2^ = 0.06), confirming that functional loss is largely independent of the common driver landscape and requires a transcriptional rather than genomic readout (Figure 3d). Two associations were nonetheless strong enough within their tumor type to be clinically useful: in colorectal cancer, BRAF mutation marked a 25% functional-loss rate versus 8% in BRAF-WT tumors (OR 3.89, 95% CI 1.95–7.89, p = 0.0004; 77% V600), and in breast cancer, HER2/ERBB2 amplification marked 21% versus 9% (OR 2.61, 95% CI 1.35–5.27, p = 0.010) (Figure 3e). These biomarker-defined subgroups (BRAF-mutant colorectal and HER2-amplified breast cancer) are enriched for functional LKB1 loss and could be prioritized for signature-based screening.

### An NRF2 co-mutation axis and an immune-cold phenotype in lung adenocarcinoma

In LUAD, signature positivity in structurally-WT tumors rose monotonically across a KRAS/NRF2 co-mutation gradient: 6% in KRAS/NRF2-pathway/LKB1-wild-type tumors, 15% with KRAS mutation alone, 35% with NRF2-pathway mutation alone (KEAP1 or NFE2L2), and 64% in KRAS/NRF2 co-mutants (Spearman trend p < 10^-10^) (Figure 4a–c). Decomposing the KEAP1/NRF2-pathway co-occurrence by LKB1 status showed it is a genuine property of the functional-loss population, not an artifact of the mutants: NRF2-pathway mutation reached 55% in functional-loss WT tumors versus 14% in signature-negative WT tumors (OR 7.7, 95% CI 4.4–13.4), comparable to the 36% seen in genomic-loss tumors (Figure 4d).

**Figure 4.**
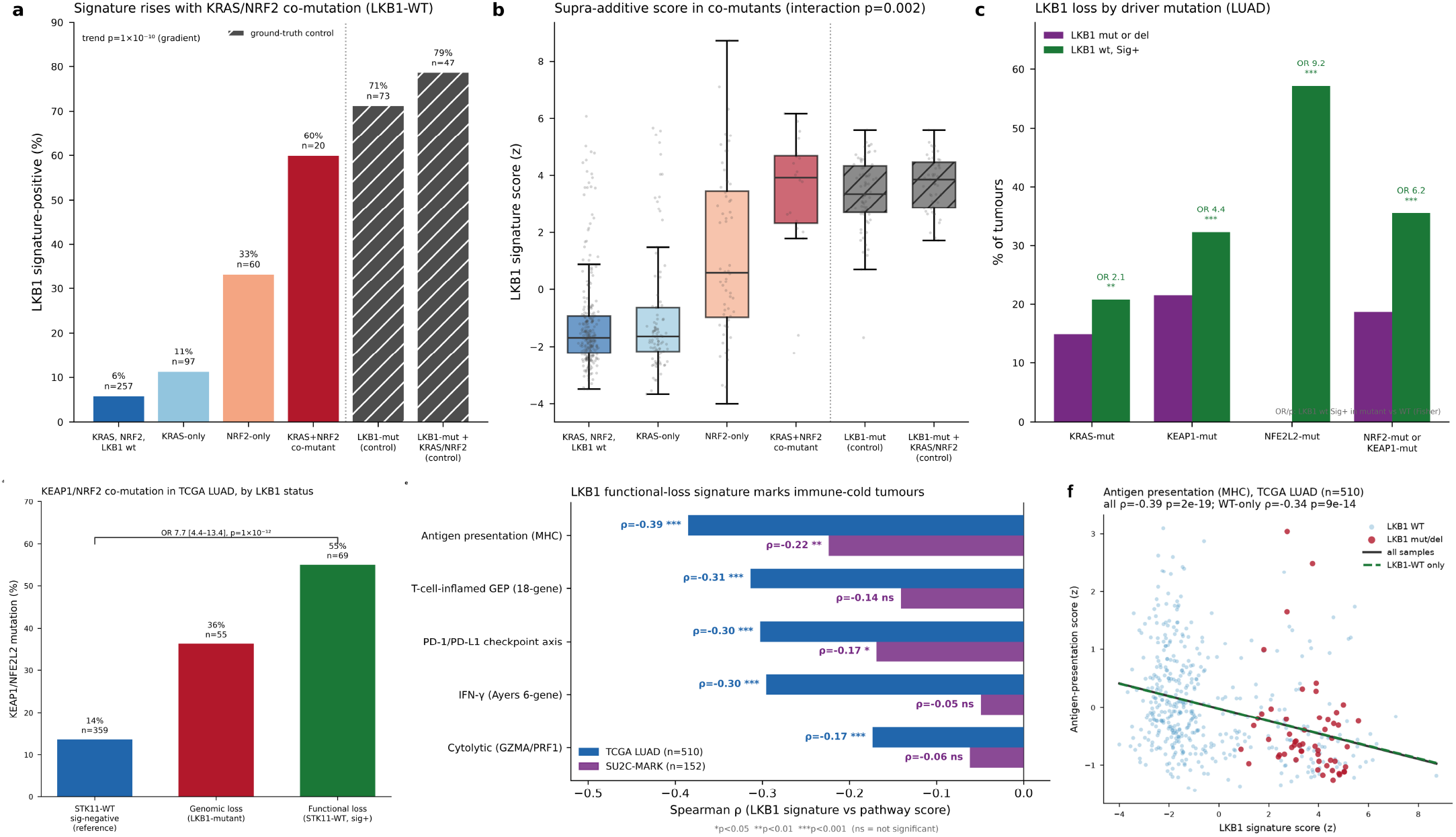
An NRF2 co-mutation axis and an immune-cold phenotype in lung adenocarcinoma. (a–c) Signature positivity in structurally wild-type LUAD rises across a KRAS/NRF2 co-mutation gradient (6% triple-wild-type → 15% KRAS-mutant → 35% NRF2-pathway-mutant [KEAP1 or NFE2L2] → 64% KRAS/NRF2 co-mutant; trend p<10?^-10^). (d) KEAP1/NRF2-pathway mutation decomposed by LKB1 status: enrichment in the functional-loss population (55%) versus signature-negative wild-type (14%; OR 7.7), comparable to genomic-loss tumors (36%). (e) The signature correlates negatively with immune programs (antigen presentation, IFN-γ, T-cell-inflamed GEP, PD-1/PD-L1 axis, cytolytic activity) in TCGA LUAD (n = 510) and SU2C-MARK (n = 152). (f) Antigen-presentation versus signature score in TCGA LUAD; the negative trend holds within genomically wild-type tumors alone (ρ = −0.34).

Consistent with this NRF2 dependence, signature-high DepMap cell lines were selectively reliant on NFE2L2 (NRF2) and its coactivator CRTC2 (a direct LKB1 target), nominating the NRF2 axis as a candidate vulnerability of the functional-loss population (Figure S4). Signature-high LUAD tumors also carried a heavier tobacco mutational imprint (SBS4 exposure ρ = +0.19; higher in genomic-loss tumors, p = 5×10^?4^), linking functional loss to the dominant LUAD mutational process (Figure S5).

Consistent with the established suppression of infiltrating immune cells in KRAS/LKB1 co-mutant (KL) lung cancer [16,20], signature-high tumors were immunologically cold. The signature correlated negatively with every immune program tested in TCGA LUAD (n = 510): antigen presentation ρ = −0.39, T-cell-inflamed GEP −0.31, PD-1/PD-L1 axis −0.30, IFN-γ −0.30, cytolytic activity −0.17 (all p < 10^?4^), and the association replicated in SU2C-MARK (n = 152; antigen presentation ρ = −0.22, p = 0.006; PD-1/PD-L1 −0.17, p = 0.037) (Figure 4e). The relationship held within genomically-WT tumors alone (TCGA antigen presentation ρ = −0.34, n = 455), showing the cold phenotype is a property of functional loss and not merely of the LKB1-mutant tumors it accompanies (Figure 4f); the inverse relationship was approximately linear across the full signature range in both cohorts (Figure S6).

## Discussion

We set out to estimate how many human tumors carry loss of LKB1 function, counting not only genomic events but also the non-genomic mechanisms that mutation-based ascertainment misses. Four findings emerge. First, a transcriptional signature of LKB1 loss can be derived rigorously from genomically-defined cases and generalizes across tumor types, independent cohorts, cell lines and platforms. Second, the signature tracks LKB1 kinase function causally: restoring wild-type (but not kinase-dead) LKB1 reverses it, the property required of a biomarker for a pathway-restoration therapy. Third, the signature reveals a functionally-deficient, structurally-intact population roughly 3.7-fold larger than the mutation-defined population, raising total LKB1-loss prevalence to ~9.9% across the 12 signature-validated cohorts (5,809 tumors) and concentrated in esophageal, colorectal, endometrial, melanoma, gastric and breast cancers where STK11 mutation is uncommon. Fourth, in lung adenocarcinoma this functional-loss population sits on a KRAS/NRF2 co-mutation axis and carries the immune-cold phenotype classically attributed to LKB1-mutant disease.

The signature is methodologically conservative: training on genomically-defined loss provides an objective label, restricting controls to confidently-intact tumors prevents contamination of the negative class, and nested cross-validation keeps discrimination unbiased. Its independent rediscovery of the LKB1 substrate SIK1, its concordance with the Kaufman classifier despite outperforming it, and its retained discrimination with STK11 removed all argue that it measures a downstream functional program. The add-back experiment converts this from correlation to causation, the only one of these lines of evidence that establishes the signature responds to LKB1 activity itself.

The genomic-correlate and NRF2-axis analyses reinforce the case for a transcriptional readout. Common drivers explained little of functional loss (pseudo-R^2^ = 0.06), yet in lung adenocarcinoma the functional-loss population is strongly enriched for NRF2-pathway (KEAP1/NFE2L2) mutation across the co-mutation gradient, recapitulating the known cooperation between LKB1 loss and NRF2 activation in KRAS-mutant lung cancer [19], and this enrichment is a property of the functional-loss population itself, not merely the LKB1 mutants. Outside lung cancer, the enrichment of functional loss in BRAF-mutant colorectal and HER2-amplified breast cancer, both defined by biomarkers already measured for every patient, offers a practical entry point for concentrating signature-based testing within routine diagnostic workflows.

The immune-cold association has a specific clinical corollary. LKB1 loss is linked to primary resistance to PD-1 blockade in KRAS-mutant lung cancer [20,21], and our signature marks immunologically cold tumors across two cohorts. However, in the SU2C-MARK immunotherapy cohort the signature did not predict checkpoint-blockade response, progression-free or overall survival, whereas STK11 mutation itself was associated with worse progression-free survival (HR 1.50), a dissociation between mutation and functional signature within a single cohort. We therefore frame the signature as identifying a cold-tumor state that may require the immune microenvironment to be changed, rather than as a checkpoint-response biomarker.

Several limitations bound the interpretation. Prevalence estimates depend on the signature-positive threshold; we anchor it to a fixed false-positive floor and report excess positivity, so absolute values are estimates with the intervals given in the supplementary tables. Within-tissue validation was strongest where genomic-loss cases were plentiful (lung, gastric) and weakest where they were few (ovarian), so low-anchor cohorts are exploratory. The analysis is confined to TCGA primary tumors and bulk expression; extension to single-cell and spatial data would refine the immune-context findings. Finally, functional loss inferred from a signature is a molecular classification; the add-back experiment provides causal support in two lines, but broader functional confirmation across tissues would strengthen it.

In sum, by grounding a transcriptional signature in genomically-defined LKB1 loss, validating it across cohorts and cell lines, and confirming that it responds to LKB1 kinase activity, we provide a mechanism-agnostic, pan-cancer estimate of LKB1 functional deficiency. The functionally-deficient but structurally-intact population is several-fold larger than the mutation-defined population and concentrated in a definable set of cancer types: a quantitative basis for any strategy that depends on restoring LKB1 function.

## Acknowledgements

The authors acknowledge the assistance of Claude Science in the research and generation of this manuscript. The results have been reviewed by the authors for accuracy.

## Supplementary items

**Fig S1**. The 30-gene LKB1 functional-loss signature: per-gene elastic-net weights (up/down in loss), effect sizes and LKB1-pathway members.

**Fig S2**. STK11 promoter methylation is uninformative in TCGA (probe distributions; no methylation–expression anticorrelation).

**Fig S3**. CCLE per-lineage STK11-loss prevalence recapitulates the TCGA tumor ranking.

**Fig S4**. DepMap dependency landscape of signature-high lines (NFE2L2/CRTC2/TYMS enrichment; no drug response survived multiple-testing correction, data not shown).

**Fig S5**. SBS4 (tobacco) mutational-signature association with the LKB1-loss signature in LUAD.

**Fig S6**. Full immune-pathway scatter grid (5 pathways × TCGA LUAD and SU2C-MARK; all-samples versus LKB1-WT-only regression).

**Note S1**. Class-imbalance robustness of signature derivation.

**Note S2**. TCGA survival associations (genomic OS HR 1.48; functional OS HR 1.31) and SU2C-MARK immunotherapy analysis (signature not predictive of response/PFS/OS; STK11 mutation associated with worse PFS, HR 1.50).

**Table S18**. The 30 signature genes with elastic-net weights, direction, univariate effect size (Δz), and Benjamini-Hochberg-adjusted Mann-Whitney p-values in TCGA LUAD.

**Supplementary Tables S1–S17** as previously enumerated, plus external- and cell-line-validation and add-back result tables.

## Supplementary figures

**Figure S1.**
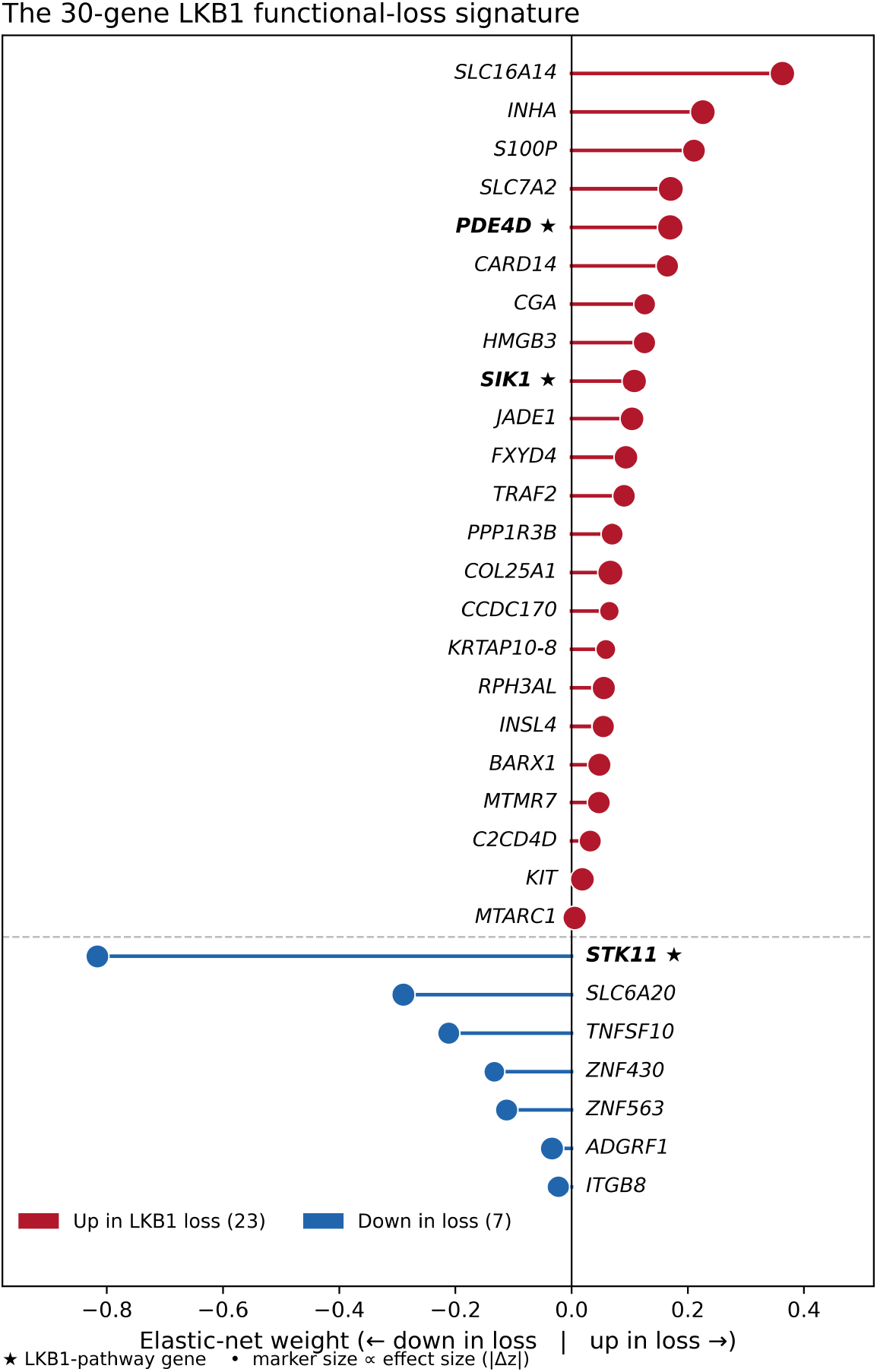
The 30-gene LKB1 functional-loss signature. Elastic-net weights of the 30 signature genes, ordered as genes up-regulated in LKB1 loss (top, 23 genes) then down-regulated genes (bottom, 7 genes), each block sorted by weight magnitude. Bar and marker color denote direction (red, up in loss; blue, down); marker size is proportional to the univariate effect size (absolute mean z-score difference between genomic-loss and wild-type LUAD tumors). Stars mark LKB1-pathway members: LKB1/STK11 itself and its downstream effectors SIK1 (a direct LKB1 substrate) and PDE4D, all recovered by the model without supervision. Per-gene weights, effect sizes and Benjamini-Hochberg-adjusted Mann-Whitney p-values (all measurable genes FDR < 4×10^?4^) are provided in Table S18.

**Figure S2.**
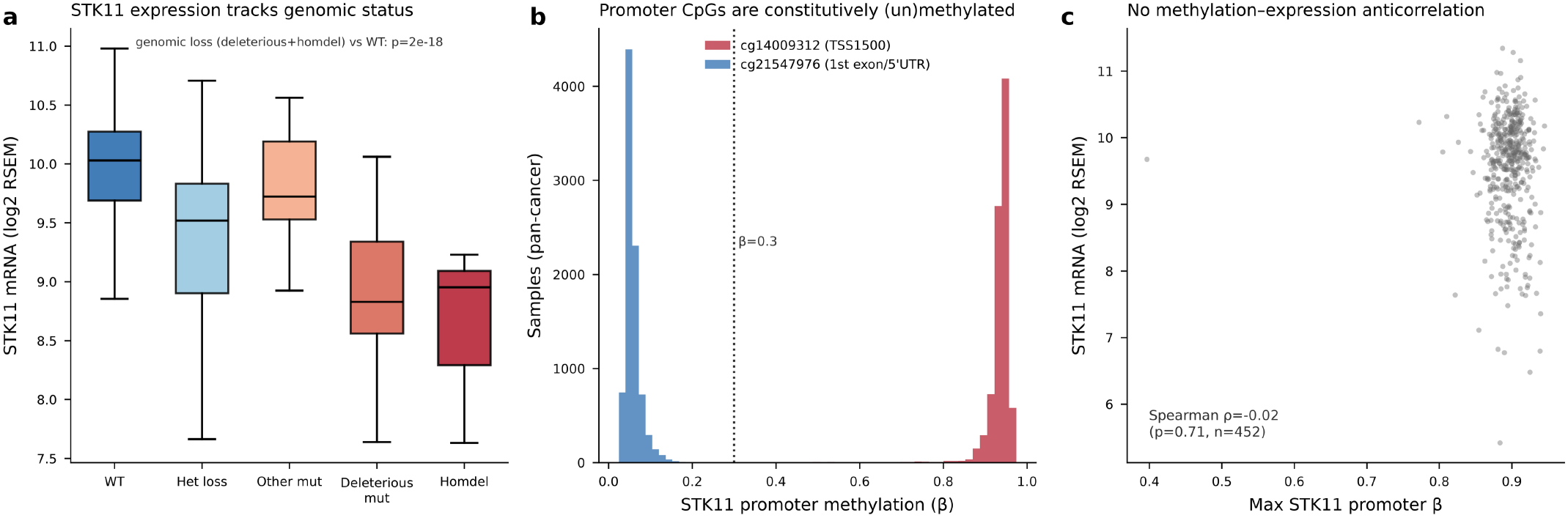
STK11 promoter methylation is not the mechanism of functional loss. (a) STK11 mRNA (log2 RSEM) by genomic status in LUAD; expression tracks genomic status (deleterious/homozygous-deletion vs wild-type, p = 2×10^-18^), confirming an expression-based readout is justified. (b) Pan-cancer distribution of the two STK11 promoter CpG probes (cg14009312, TSS1500; cg21547976, 1st exon/5′UTR): both are constitutively (un)methylated (one near β≈0.9, the other near β≈0.05), with essentially no intermediate values. (c) No anticorrelation between maximum promoter methylation and STK11 expression (Spearman ρ = −0.02, p = 0.71, n = 452), ruling out promoter hypermethylation as a pan-cancer silencing mechanism.

**Figure S3.**
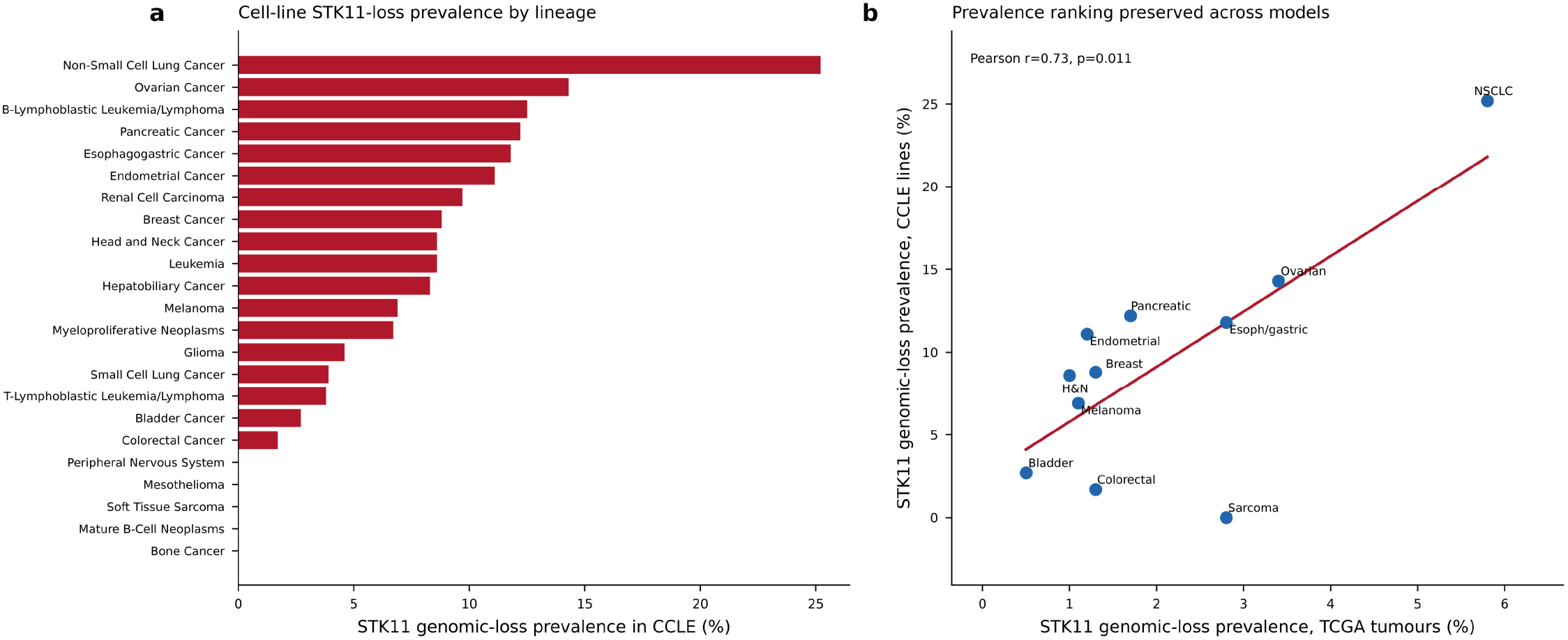
Cell-line STK11-loss prevalence recapitulates the tumor ranking. (a) STK11 genomic-loss prevalence by lineage across CCLE cell lines, led by non-small-cell lung cancer (~25%). (b) Per-lineage STK11 genomic-loss prevalence in CCLE lines versus matched TCGA tumor types; the ranking is preserved (Pearson r = 0.73, p = 0.011), while cell lines systematically over-represent STK11 loss relative to primary tumors.

**Figure S4.**
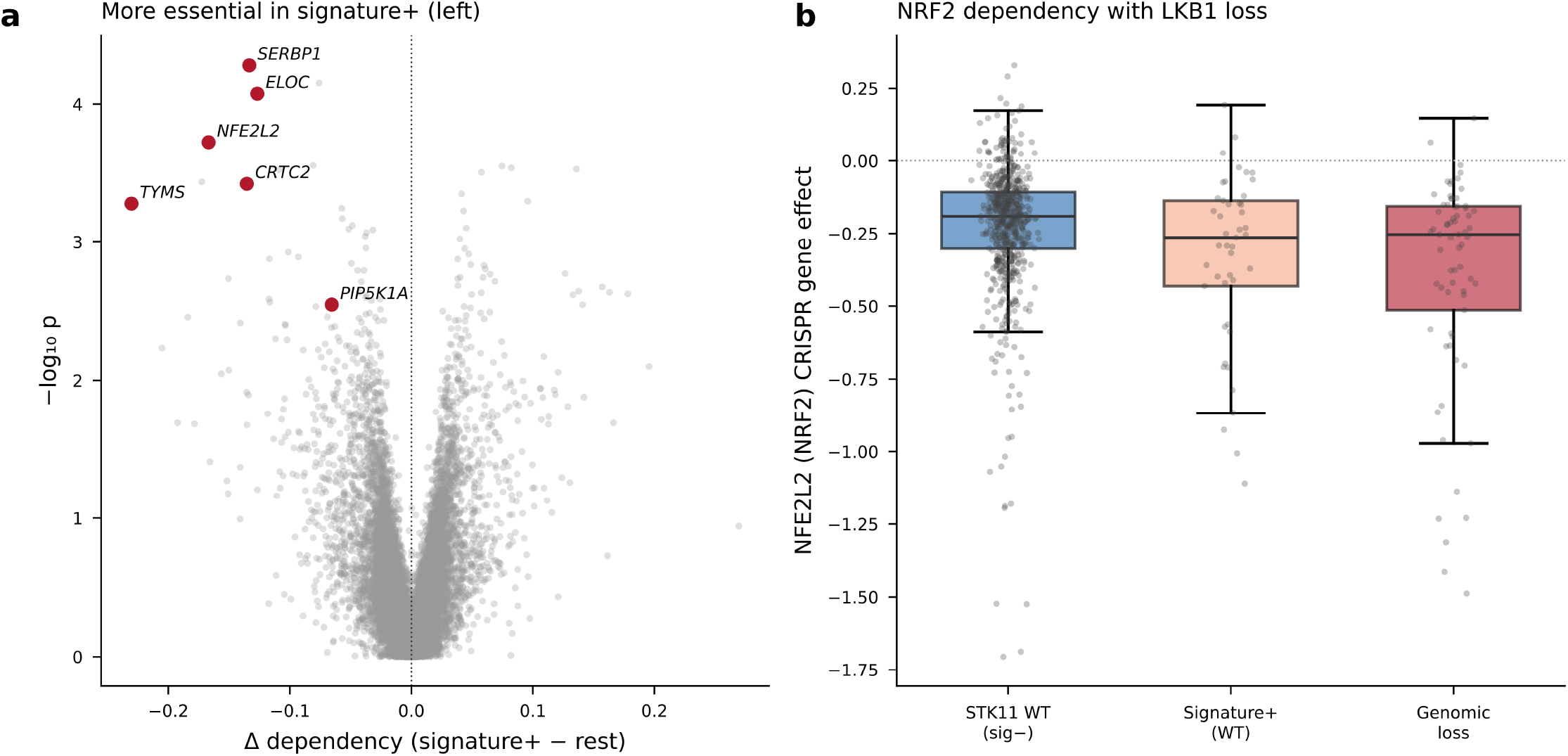
DepMap dependency landscape of signature-high cell lines. (a) Volcano of the difference in CRISPR gene effect between signature-positive lines and the rest (x-axis, more negative = more essential in signature-positive; y-axis, −log10 p). NFE2L2 (NRF2), its partner CRTC2, the thymidylate synthase TYMS, and translation/elongation factors (SERBP1, ELOC) are among the top selective dependencies of signature-high lines. (b) NFE2L2 (NRF2) CRISPR gene effect by group: STK11 wild-type/signature-negative, signature-positive wild-type, and genomic loss, showing progressively greater NRF2 dependence with LKB1 loss. No small-molecule sensitivity survived multiple-testing correction (data not shown).

**Figure S5.**
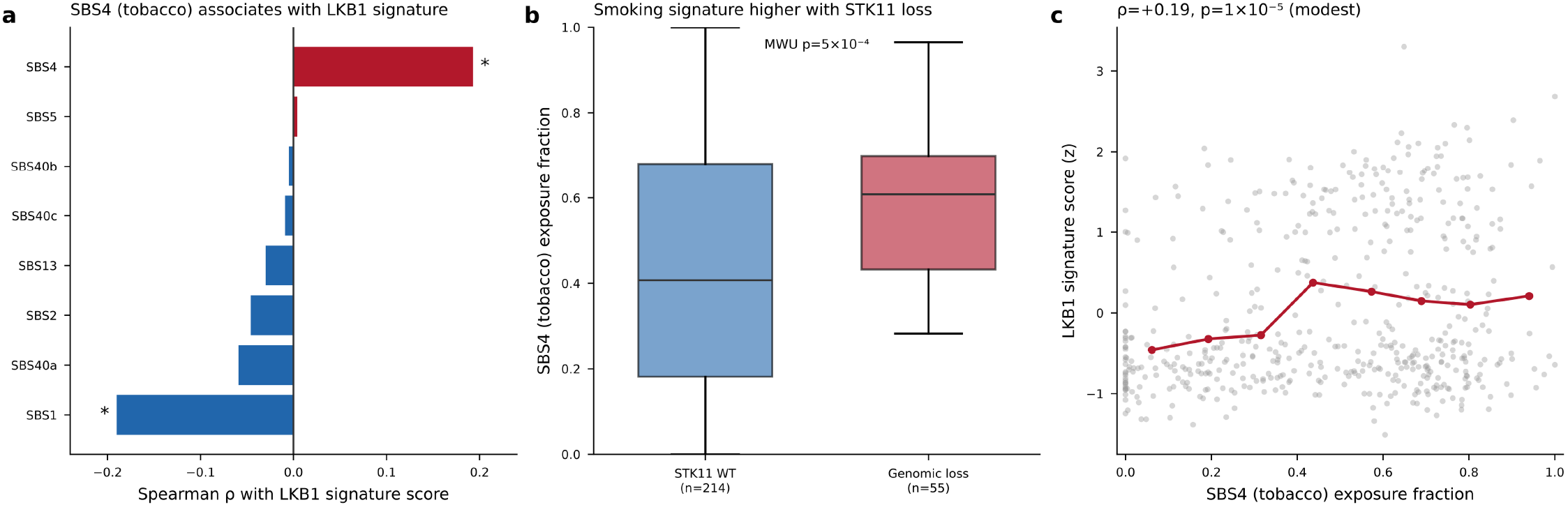
The tobacco mutational signature SBS4 associates with LKB1 functional loss in LUAD. (a) Spearman correlation of each COSMIC SBS exposure with the LKB1 signature score; SBS4 (tobacco) is the strongest positive association and SBS1 (clock-like) the strongest negative (* marks FDR-significant). (b) SBS4 exposure fraction is higher in genomic-loss than in STK11 wild-type tumors (Mann-Whitney p = 5×10^?4^). (c) LKB1 signature score versus SBS4 exposure fraction (Spearman ρ = +0.19, p = 1×10^?5^); the association is real but modest, indicating smoking-related mutagenesis is a contributor but not the sole driver of functional loss.

**Figure S6.**
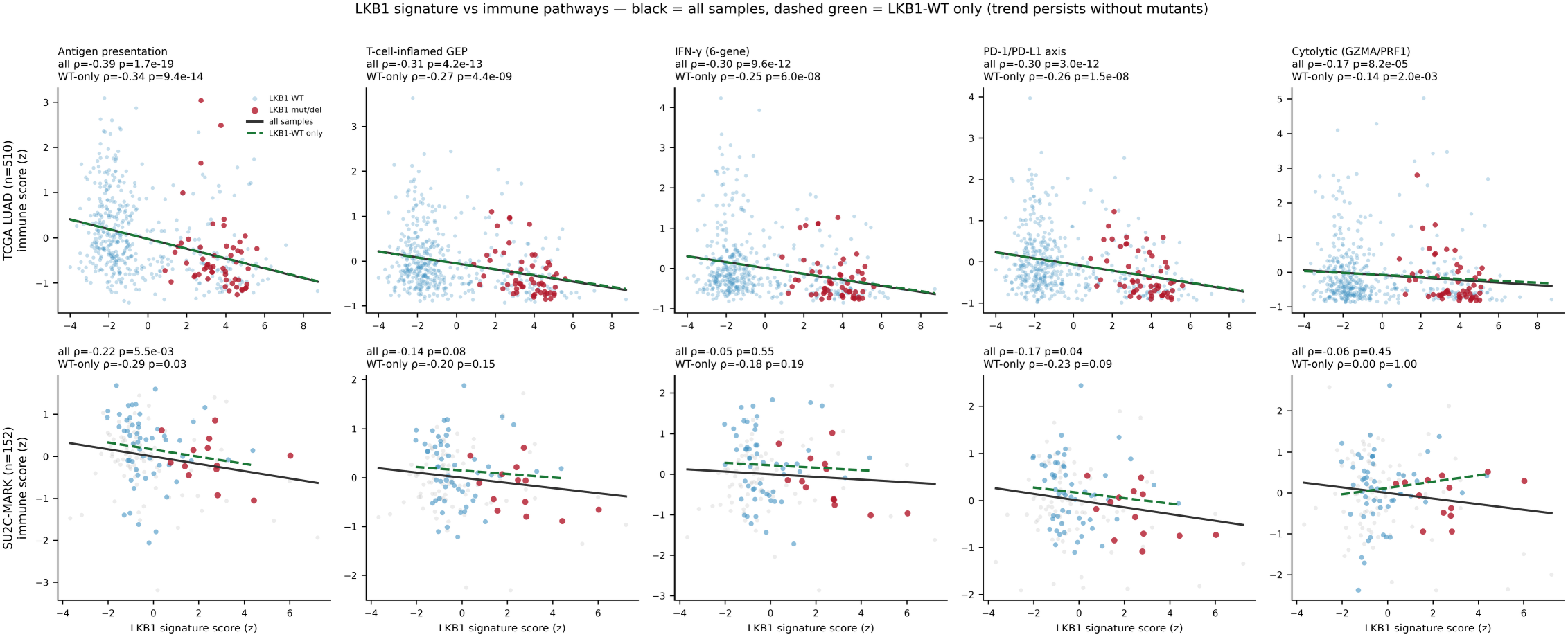
The LKB1 signature is inversely related to immune-pathway activity across two cohorts. Scatter of five immune-pathway scores (columns: antigen presentation, T-cell-inflamed GEP, IFN-γ 6-gene, PD-1/PD-L1 axis, cytolytic GZMA/PRF1) against the LKB1 functional-loss signature score in TCGA LUAD (top row, n = 510) and SU2C-MARK (bottom row, n = 152). LKB1 structurally wild-type tumors in blue, LKB1 mutant/deleted tumors in red. The solid black line is the regression across all samples; the dashed green line is the regression restricted to LKB1 wild-type tumors, showing the inverse relationship persists after excluding genomically-altered cases. Spearman ρ and p-values (all-samples and WT-only) are annotated per panel.

## References

1. A serine/threonine kinase gene defective in Peutz-Jeghers syndrome. Nature. 1998. doi:10.1038/34432

2. Peutz-Jeghers syndrome is caused by mutations in a novel serine threonine kinase. Nature Genetics. 1998. doi:10.1038/ng0198-38

3. Peutz-Jeghers syndrome. Current Opinion in Gastroenterology. 2021. doi:10.1097/mog.0000000000000718

4. MO25α/β interact with STRADα/β enhancing their ability to bind, activate and localize LKB1 in the cytoplasm. The EMBO Journal. 2003. doi:10.1093/emboj/cdg490

5. Complexes between the LKB1 tumor suppressor, STRADα/β and MO25α/β are upstream kinases in the AMP-activated protein kinase cascade. Journal of Biology. 2003. doi:10.1186/1475-4924-2-28

6. LKB1 Is the Upstream Kinase in the AMP-Activated Protein Kinase Cascade. Current Biology. 2003. doi:10.1016/j.cub.2003.10.031

7. The tumor suppressor LKB1 kinase directly activates AMP-activated kinase and regulates apoptosis in response to energy stress. PNAS. 2004. doi:10.1073/pnas.0308061100

8. The LKB1 tumor suppressor negatively regulates mTOR signaling. Cancer Cell. 2004. doi:10.1016/j.ccr.2004.06.007

9. LKB1 is a master kinase that activates 13 kinases of the AMPK subfamily, including MARK/PAR-1. The EMBO Journal. 2004. doi:10.1038/sj.emboj.7600110

10. LKB1-Dependent Signaling Pathways. Annual Review of Biochemistry. 2006. doi:10.1146/annurev.biochem.75.103004.142702

11. LKB1; linking cell structure and tumor suppression. Oncogene. 2008. doi:10.1038/onc.2008.342

12. The LKB1-AMPK pathway: metabolism and growth control in tumor suppression. Nature Reviews Cancer. 2009. doi:10.1038/nrc2676

13. LKB1 biology: assessing the therapeutic relevancy of LKB1 inhibitors. Cell Communication and Signaling. 2024. doi:10.1186/s12964-024-01689-5

14. Prevalence and specificity of LKB1 genetic alterations in lung cancers. Oncogene. 2007. doi:10.1038/sj.onc.1210418

15. LKB1 modulates lung cancer differentiation and metastasis. Nature. 2007. doi:10.1038/nature06030

16. Co-occurring Genomic Alterations Define Major Subsets of KRAS-Mutant Lung Adenocarcinoma with Distinct Biology, Immune Profiles, and Therapeutic Vulnerabilities. Cancer Discovery. 2015. doi:10.1158/2159-8290.cd-14-1236

17. STK11/LKB1 Deficiency Promotes Neutrophil Recruitment and Proinflammatory Cytokine Production to Suppress T-cell Activity in the Lung Tumor Microenvironment. Cancer Research. 2016. doi:10.1158/0008-5472.can-15-1439

18. Suppression of STING Associated with LKB1 Loss in KRAS-Driven Lung Cancer. Cancer Discovery. 2018. doi:10.1158/2159-8290.cd-18-0689

19. LKB1 and KEAP1/NRF2 Pathways Cooperatively Promote Metabolic Reprogramming with Enhanced Glutamine Dependence in KRAS-Mutant Lung Adenocarcinoma. Cancer Research. 2019. doi:10.1158/0008-5472.can-18-3527

20. STK11/LKB1 Mutations and PD-1 Inhibitor Resistance in KRAS-Mutant Lung Adenocarcinoma. Cancer Discovery. 2018. doi:10.1158/2159-8290.cd-18-0099

21. Co-occurring genomic alterations in non-small-cell lung cancer biology and therapy. Nature Reviews Cancer. 2019. doi:10.1038/s41568-019-0179-8

22. Targeting LKB1 in cancer - exposing and exploiting vulnerabilities. British Journal of Cancer. 2015. doi:10.1038/bjc.2015.261

23. Epigenetic inactivation of LKB1 in primary tumors associated with the Peutz-Jeghers syndrome. Oncogene. 2000. doi:10.1038/sj.onc.1203227

24. 5′-CpG island methylation of the LKB1/STK11 promoter and allelic loss at chromosome 19p13.3 in sporadic colorectal cancer. Gut. 2000. doi:10.1136/gut.47.2.272

25. STK11/LKB1 Loss of Function Is Associated with Global DNA Hypomethylation and S-Adenosyl-Methionine Depletion in Human Lung Adenocarcinoma. Cancer Research. 2021. doi:10.1158/0008-5472.can-20-3199

26. LKB1 loss links serine metabolism to DNA methylation and tumorigenesis. Nature. 2016. doi:10.1038/nature20132

27. LKB1 Loss at Transcriptional Level Promotes Tumor Malignancy and Poor Patient Outcomes in Colorectal Cancer. Annals of Surgical Oncology. 2014. doi:10.1245/s10434-014-3824-1

28. LKB1 Down-Modulation by miR-17 Identifies Patients With NSCLC Having Worse Prognosis Eligible for Energy-Stress-Based Treatments. Journal of Thoracic Oncology. 2021. doi:10.1016/j.jtho.2021.04.005

29. Negative Regulation of Serine Threonine Kinase 11 (STK11) through miR-100 in Head and Neck Cancer. Genes. 2020. doi:10.3390/genes11091058

30. Post-translational regulation contributes to the loss of LKB1 expression through SIRT1 deacetylase in osteosarcomas. British Journal of Cancer. 2017. doi:10.1038/bjc.2017.174

31. Posttranslational regulation of liver kinase B1 in human cancer. Journal of Biological Chemistry. 2023. doi:10.1016/j.jbc.2023.104570

32. Immunohistochemical Loss of LKB1 Is a Biomarker for More Aggressive Biology in KRAS-Mutant Lung Adenocarcinoma. Clinical Cancer Research. 2015. doi:10.1158/1078-0432.ccr-14-3112

33. Somatic LKB1 Mutations Promote Cervical Cancer Progression. PLoS ONE. 2009. doi:10.1371/journal.pone.0005137

34. STK11/LKB1 Peutz-Jeghers Gene Inactivation in Intraductal Papillary-Mucinous Neoplasms of the Pancreas. American Journal of Pathology. 2001. doi:10.1016/s0002-9440(10)63053-2

35. LKB1 Haploinsufficiency Cooperates With Kras to Promote Pancreatic Cancer Through Suppression of p21-Dependent Growth Arrest. Gastroenterology. 2010. doi:10.1053/j.gastro.2010.04.055

36. Loss of Stk11/Lkb1 Expression in Pancreatic and Biliary Neoplasms. Modern Pathology. 2003. doi:10.1097/01.mp.0000075645.97329.86

37. The role of STK11/LKB1 in cancer biology: implications for ovarian tumorigenesis and progression. Frontiers in Cell and Developmental Biology. 2024. doi:10.3389/fcell.2024.1449543

38. LKB1 Loss Induces Characteristic Patterns of Gene Expression in Human Tumors Associated with NRF2 Activation and Attenuation of PI3K-AKT. Journal of Thoracic Oncology. 2014. doi:10.1097/jto.0000000000000173

39. Integrative Genomic and Proteomic Analyses Identify Targets for Lkb1-Deficient Metastatic Lung Tumors. Cancer Cell. 2010. doi:10.1016/j.ccr.2010.04.026

40. Genomic landscape of lung adenocarcinoma in East Asians. Nature Genetics. 2020. doi:10.1038/s41588-019-0569-6

41. Proteogenomic Characterization Reveals Therapeutic Vulnerabilities in Lung Adenocarcinoma. Cell. 2020. doi:10.1016/j.cell.2020.06.013

42. Integrative Proteomic Characterization of Human Lung Adenocarcinoma. Cell. 2020. doi:10.1016/j.cell.2020.05.043

43. Tumor mutational load predicts survival after immunotherapy across multiple cancer types. Nature Genetics. 2019. doi:10.1038/s41588-018-0312-8

44. Tumor and Microenvironment Evolution during Immunotherapy with Nivolumab. Cell. 2017. doi:10.1016/j.cell.2017.09.028

45. Distinct Immune Cell Populations Define Response to Anti-PD-1 Monotherapy and Anti-PD-1/Anti-CTLA-4 Combined Therapy. Cancer Cell. 2019. doi:10.1016/j.ccell.2019.01.003

46. Genomic and transcriptomic determinants of response to immune checkpoint blockade in NSCLC (SU2C-MARK). Nature Genetics. 2023. doi:10.1038/s41588-023-01355-5

47. Next-generation characterization of the Cancer Cell Line Encyclopedia. Nature. 2019. doi:10.1038/s41586-019-1186-3

48. Kretschmer LS, Mitchell DC, Liu J, et al. Small molecule activation of the tumor suppressor kinase LKB1. bioRxiv. 2024. doi:10.1101/2024.12.17.628051

